# Food seeking suppression by environmental enrichment accompanies cell type- and circuit-specific prelimbic cortical modulation

**DOI:** 10.1101/2025.06.11.659116

**Authors:** Kate Z. Peters, Romarua Agbude, Oliver G. Steele, Nobuyoshi Suto, Eisuke Koya

## Abstract

Cues such as fast-food advertisements associated with food can provoke food cravings which may lead to unhealthy overeating. To effectively control such cravings, we need to better understand the factors that reduce food cue reactivity and reveal corresponding ‘anti-craving’ brain mechanisms. We previously reported that access to environmental enrichment (EE) that provides cognitive and physical stimulation in mice reduced cue-evoked sucrose seeking and prelimbic cortex (PL) neuronal reactivity.

To date, the phenotype of PL neurons that undergo EE-induced adaptations has not been fully elucidated. Therefore, we used brain slice electrophysiology to investigate how EE modulated intrinsic excitability in the general population of PL interneurons and pyramidal cells. Additionally, we used retrograde tracing and the neuronal activity marker ‘Fos’ to investigate how EE modulated cue-evoked recruitment of pyramidal cells projecting to the paraventricular nucleus of the thalamus (PVT) and nucleus accumbens core (NAc).

Before the cue-evoked sucrose seeking test, EE enhanced the general, baseline excitability of inhibitory interneurons, but not pyramidal cells, thereby promoting inhibitory ‘overdrive’. During cue-evoked sucrose seeking, EE suppressed recruitment of PVT-, but not NAc-projecting, neurons thereby selectively promoting corticothalamic, but not corticoaccumbens, excitatory ‘underdrive’. Collectively, we further illuminate EE’s ‘anti-food seeking’ actions whereby EE promotes both cell type-specific (inhibitory interneuron ‘overdrive’) and circuit-specific (excitatory corticothalamic ‘underdrive’) neuroadaptations in the PL.

## Introduction

Signals or ‘cues’ associated with palatable foods (e.g. fast-food advertisements) trigger food memory retrieval, but also responses such as food cravings that trigger unhealthy overeating. In laboratory animals, these cues invigorate food seeking behavior, allowing us to conduct detailed neurobiological investigations into food cue reactivity. To date, such research has largely been performed to reveal ‘pro-craving’ brain circuits that provoke cue reactivity, but there is a paucity of information about ‘anti-craving’ circuits that suppress cue reactivity.

We and others have provided physical and cognitive stimulation to laboratory rodents by providing housing with environmental enrichment (EE) through provision of larger cages with exercise wheels, tunnels and shelters, toys, and multiple nesting materials (Nithianantharajah & Hannan, 2006; Grimm & Sauter, 2020; Solinas *et al*., 2020; Margetts-Smith *et al*., 2021) (**Fig 1a)**. One day EE exposure attenuated cue-evoked sucrose seeking in both mice and rats (Grimm & Sauter, 2020; Margetts-Smith *et al*., 2021). Furthermore, EE attenuated the recruitment of sparse sets of activated neurons or ‘neuronal ensembles’ expressing the neuronal activity marker ‘Fos’ in the prelimbic area (PL) of the medial prefrontal cortex (mPFC), a brain structure that controls motivated actions (Dalley *et al*., 2004; Riga *et al*., 2014; Gourley & Taylor, 2016). Specifically, EE reduced Fos in PL’s deep layer neurons, which project to various subcortical, thalamic and striatal structures that coordinate appetitive behaviors (Berendse *et al*., 1992; Voorn *et al*., 2004; Vertes, 2006; Riga *et al*., 2014; Anastasiades & Carter, 2021).

**Figure 1.**
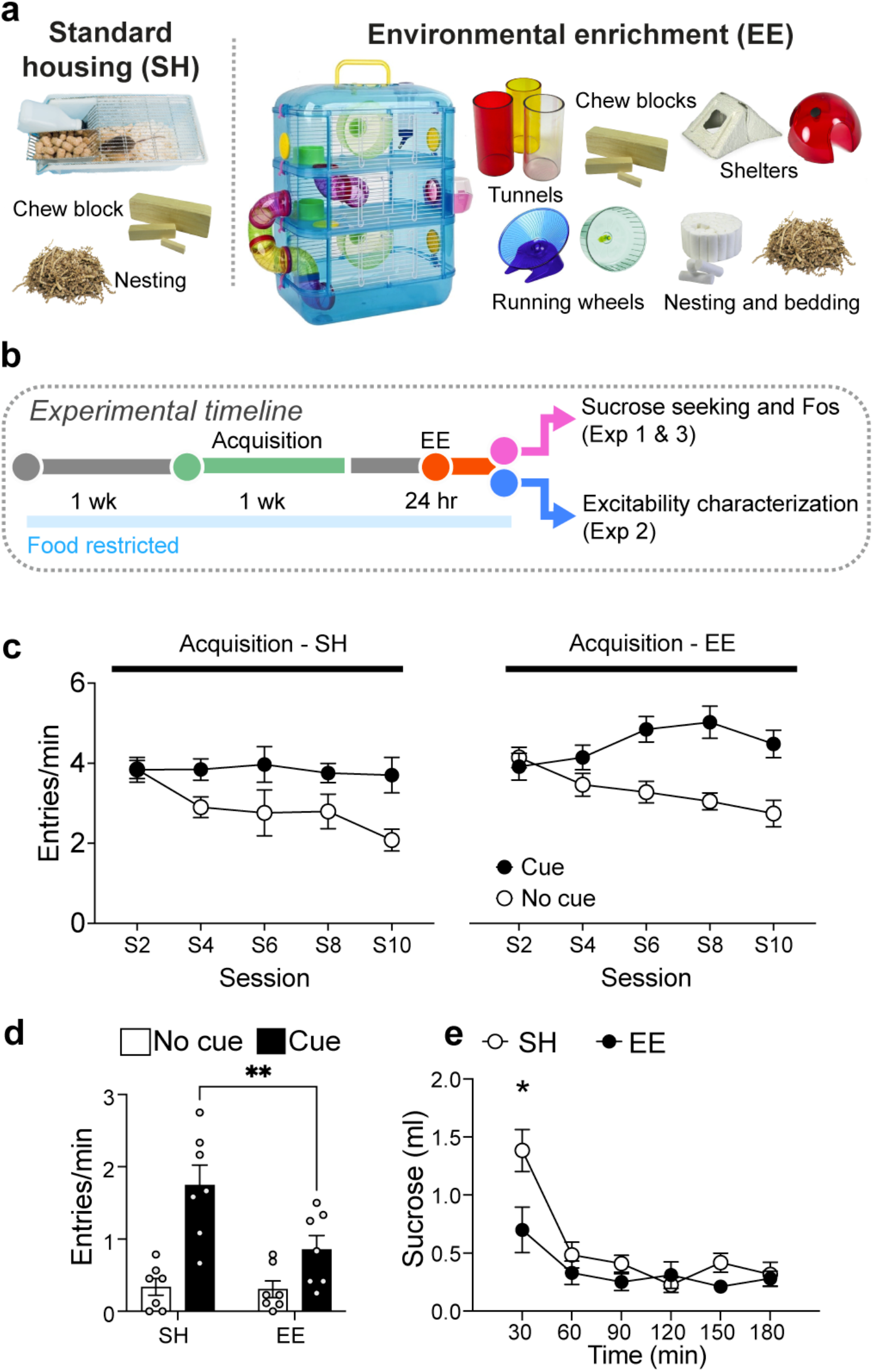
Experimental timelines and housing conditions. **a** Images of housing with environmental enrichment (EE) and control standard housing (SH), **b** Timeline. **c** Head entry rates into the magazine during the Cue and No Cue periods during acquisition for mice in the EE and SH conditions (n=7/group). **d** EE selectively modulates cue-evoked sucrose seeking. **e** EE attenuates sucrose consumption (n=6-7/group). \*\**p* < 0.01 and \**p* < 0.05 against mice in SH condition. All data are expressed as mean±SEM.

Alterations in mPFC neuronal excitability and recruitment underlie the modulation of cue-evoked food seeking (Brebner, Ziminski, Margetts-Smith, Sieburg, Hall, *et al*., 2020; Brebner, Ziminski, Margetts-Smith, Sieburg, Reeve, *et al*., 2020). In the mPFC, both inhibitory interneurons that increase local inhibition and excitatory pyramidal cells that project to various subcortical structures that subserve motivation and reward orchestrate cue-evoked reward seeking behaviors.(Sparta *et al*., 2014; Otis *et al*., 2017; Brebner, Ziminski, Margetts-Smith, Sieburg, Hall, *et al*., 2020; Brebner, Ziminski, Margetts-Smith, Sieburg, Reeve, *et al*., 2020). Here we hypothesized that EE promoted the suppression of cue-evoked food seeking through modulating the excitability of PL neurons and recruitment of PL projection neurons to the paraventricular nucleus of the thalamus (PVT) and nucleus accumbens core (NAc), which play key roles in processing cue-related information and orchestrating reward seeking (Otis *et al*., 2017; Iglesias & Flagel, 2021). Since women experience more food cravings and exhibit higher cortical activation following food cue exposure than men (Pelchat, 1997; Zellner *et al*., 1999; Uher *et al*., 2006; Hallam *et al*., 2016), we focused our investigations on examining EE effects on modulating PL excitability and projection neuron recruitment in female mice.

## Results

### Experiment 1 – EE reduces cue-evoked food seeking and sucrose consumption

#### Pavlovian Conditioning

We conditioned mice to associate an auditory cue with the delivery of 10% sucrose solution across 12 sessions (**Fig 1b**). Mice assigned to the EE and SH conditions acquired a cue-reward association at similar levels and exhibited selective head entry responding during Cue presentation period (auditory clicker) compared to No Cue period (Session X EE x Cue interaction, *F*_(9,126)_ = 0.733, *p* = 0.6776; Session x Cue, *F*_(9,126)_ = 5.468; *p* < 0.0001; **Fig 1c**).

#### Cue-evoked sucrose seeking and sucrose consumption test

We assessed the effects of EE on cue-evoked sucrose seeking under extinction conditions. Here, sucrose conditioned mice were either exposed to 1d EE or remained in standard housing (SH controls) and underwent testing. Similar to our previous study (Margetts-Smith *et al*., 2021), EE selectively reduced cue-evoked sucrose seeking (EE x Cue, *F*_(1,12)_ = 7.273, *p* < 0.05; Main effects of EE *F*_(1,12)_ = 4.857, *p* < 0.05 and Cue *F*_(1,12)_ = 37.90, *p* < 0.001; **Fig 1d**). An additional cohort of sucrose conditioned, EE-exposed mice (or not exposed) underwent sucrose consumption testing. EE significantly attenuated sucrose consumption (EE x Time, *F*_(5,45)_ = 3.146, *p* < 0.05; Main effect of EE, *F*_(1,9)_ = 5.967, *p* < 0.05; **Fig 1e**).

### Experiment 2 – EE enhances baseline PL interneuron, but not pyramidal cell excitability

We previously reported that EE reduced cue-evoked Fos expression PL (Margetts-Smith *et al*., 2021). This finding suggests that EE modulates neuronal properties, such as intrinsic excitability that influences neuronal activity (Kourrich *et al*., 2015; Ziminski *et al*., 2017; Brebner, Ziminski, Margetts-Smith, Sieburg, Reeve, *et al*., 2020), even prior to cue exposure. Thus, we examined whether EE modulated ‘baseline’ (prior to test session) excitability in PL pyramidal cells and interneurons in layers V-VI, immediately following 1d of EE.

#### EE effects on PL pyramidal cell excitability

We analyzed the number of action potentials elicited in response to positive current step injections from pyramidal cells from mice in EE and SH conditions. EE did not modulate the baseline intrinsic excitability of PL pyramidal neurons as EE did not significantly modulate firing capacity (**Fig 2a, 2b**; EE Condition × Current; *F*_(20, 700)_ = 1.23, *p* = 0.22) and active and passive membrane properties **(Fig 2c-f; Table 1)**.

**Table 1:**
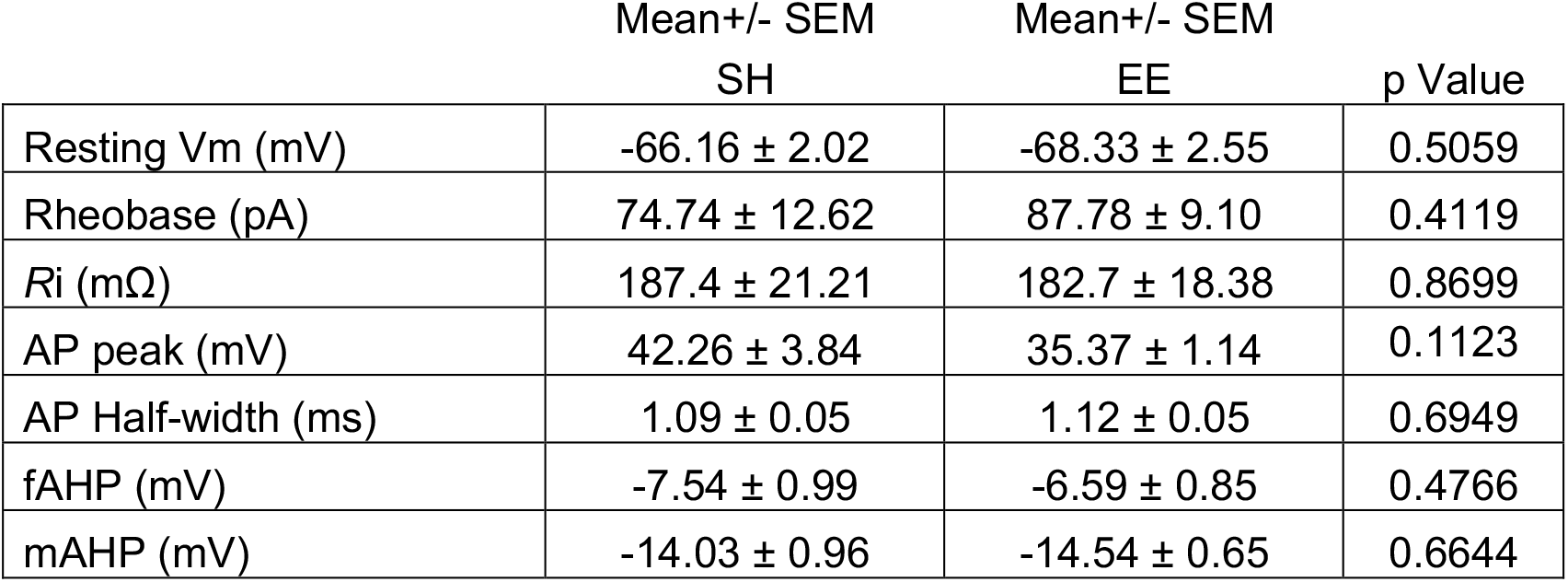
Basic membrane properties from PL pyramidal cells following EE. Basic membrane properties from PL pyramidal cells from mice in EE and SH (no EE) conditions. V_m_, Resting membrane potential; R_i_, input resistance; AP, action potential; fAHP, fast afterhyperpolarization; mAHP, medium afterhyperpolarization. All data are expressed as mean ± SEM.

|  | Mean+/- SEM<br>SH | Mean+/- SEM<br>EE | p Value |
| --- | --- | --- | --- |
| Resting $V_m$ (mV) | -66.16 ± 2.02 | -68.33 ± 2.55 | 0.5059 |
| Rheobase (pA) | 74.74 ± 12.62 | 87.78 ± 9.10 | 0.4119 |
| $R_i$ (mΩ) | 187.4 ± 21.21 | 182.7 ± 18.38 | 0.8699 |
| AP peak (mV) | 42.26 ± 3.84 | 35.37 ± 1.14 | 0.1123 |
| AP Half-width (ms) | 1.09 ± 0.05 | 1.12 ± 0.05 | 0.6949 |
| fAHP (mV) | -7.54 ± 0.99 | -6.59 ± 0.85 | 0.4766 |
| mAHP (mV) | -14.03 ± 0.96 | -14.54 ± 0.65 | 0.6644 |

**Figure 2.**
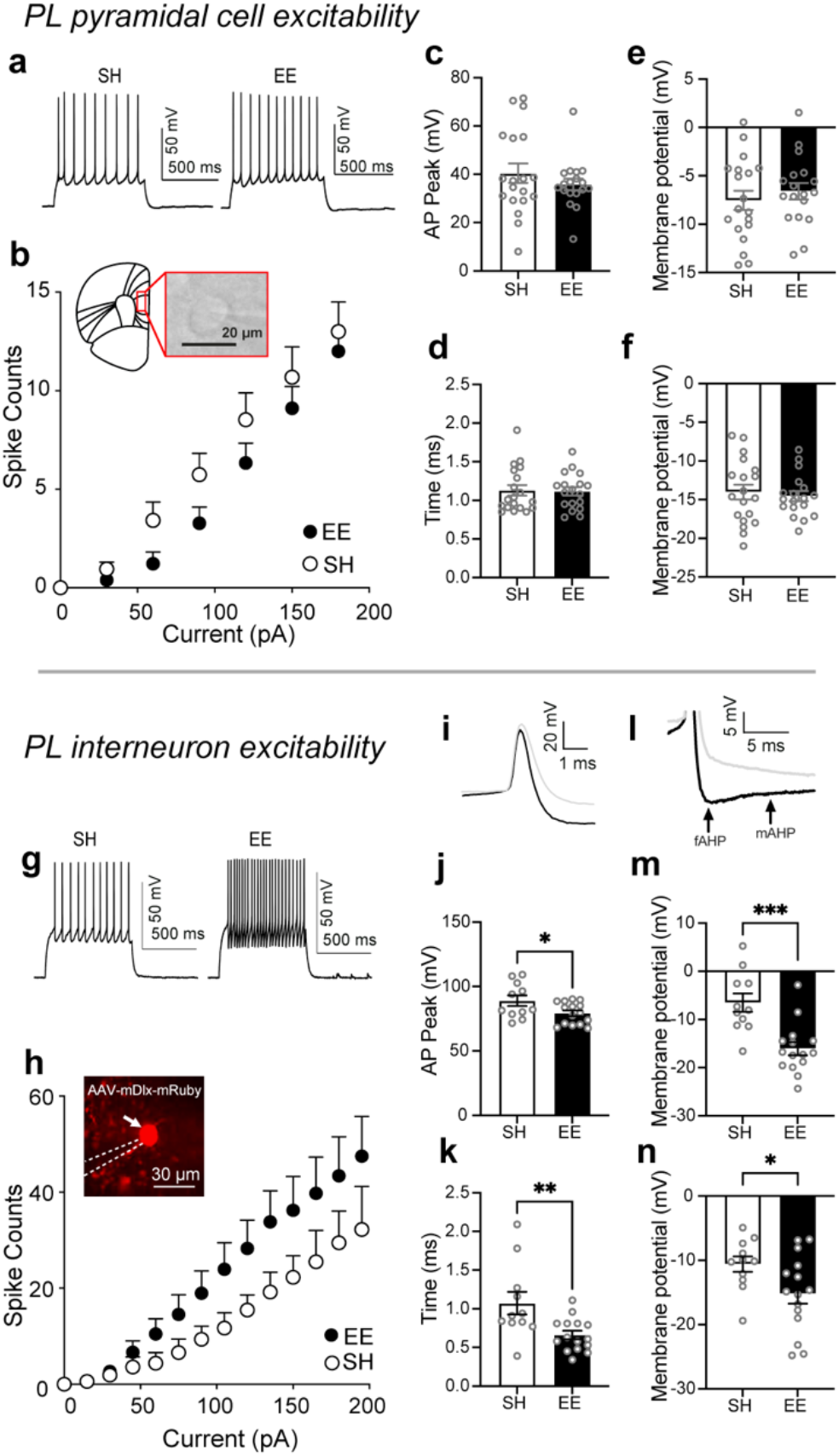
EE does not modulate the intrinsic excitability of prelimbic (PL) pyramidal cells (a-f), but of PL interneurons (g-n). **a** Representative current injection traces and **b** Firing capacity of pyramidal cells from EE or SH (no EE) conditions (SH: n=19/5, EE: n=18/5), inset: representative pyramidal cell during recording. **c** action potential (AP), peak **d** half-width, **e** fast afterhyperpolarization (fAHP), **f** medium AHP (mAHP) from pyramidal cells from EE and SH conditions. **g** Representative current injection traces and **h** firing capacity of interneurons from EE and SH conditions (SH: n=11/5, EE: n=15/6), inset: representative mRuby+ interneuron during recording (indicated by white arrow). **i** Representative action potential and **l** afterhyperpolarization (AHP) trace from interneurons from mice in EE (black line) and SH (gray line) conditions. **j** AP peak, **k** AP half-width, **m** fAHP, and **n** mAHP of interneurons from mice in EE and SH conditions. *n* = total number of cells/total number of mice. All data are expressed as mean±SEM.

#### EE effects on PL interneuron excitability

Mice were injected with AAV-mDlx-mRuby (Dimidschstein *et al*., 2016) in PL to virally-express the red fluorescent protein ‘mRuby’ in GABAergic interneurons and we measured their excitability. EE significantly upregulated firing capacity **(Fig 2g, 2h**; EE Condition × Current; F_(87, 1979)_ = 11.17, *p* < 0.001; main effect of EE (F_(1, 24)_= 5.94, *p <* 0.05) and Current (*F*_(3.18, 72.35)_ = 11.17, *p <* 0.001). In addition, we observed decreases in action potential (AP) peak (**Fig 2i and 2j;** *t*_(24)_ = 2.233, *p* < 0.05), AP half-width (**Fig 2i and 2k;** *t*_(24)_ = 2.930, *p* < 0.01), and in the fast and medium after hyperpolarization (**Fig 2l and 2m**; fAHP *t*_(24)_ = 4.221, *p* < 0.001; **Fig 2l and 2n;** mAHP *t*_(24)_ = 2.221, *p* < 0.05). No significant differences were observed for the resting membrane potential, rheobase, and input resistance (**Table 2)**. Hence, enhanced firing in the EE condition was attributable to a narrowing of the action potentials.

**Table 2:** Basic membrane properties from PL interneurons following EE. Basic membrane properties from mRuby+, PL interneurons from mice in EE and SH (no EE) conditions. V_m_, Resting membrane potential; R_i_, input resistance; AP, action potential; fAHP, fast afterhyperpolarization; mAHP, medium afterhyperpolarization. *p<0.05, **p<0.01. All data are expressed as mean ± SEM.

|  | Mean+/- SEM<br>SH | Mean+/- SEM<br>EE | p Value |
| --- | --- | --- | --- |
| Resting V<sub>m</sub> (mV) | -65.41 ± 2.20 | -69.67 ± 2.09 | 0.1815 |
| Rheobase (pA) | 64.55 ± 12.60 | 72.33 ± 16.94 | 0.7334 |
| R<sub>i</sub> (mΩ) | 287.5 ± 40.90 | 312.40 ± 27.11 | 0.6002 |
| AP peak (mV) | 88.92 ± 4.10 | 79.26 ± 2.18 * | 0.0351 |
| AP Half-width (ms) | 1.07 ± 0.15 | 0.66 ± 0.06 ** | 0.0073 |
| fAHP (mV) | -6.51 ± 1.90 | -16.08 ± 1.36 * | 0.0003 |
| mAHP (mV) | -10.59 ± 1.21 | -15.20 ± 1.54 * | 0.0361 |

### Experiment 3 – EE attenuates Fos in PL→PVT, but not PL→NAc neurons

Using two retrograde AAVs, we labeled PL neurons that project to the nucleus accumbens (NAc) and paraventricular nucleus of the thalamus (PVT) in the same mice with either GFP or mCherry. We then assessed Fos expression in these neuronal populations following EE. Viral injections sites are shown in **Figure 3a** and representative images of PL immunohistochemistry in **Figure 3b**. Retrograde labeling of PL→NAc and PL→PVT neurons was similar across mice in EE and SH (PL→NAc: (*t*_(15)_ = 0.685, *p* = 0.504), PL→PVT (*t*_(15)_ = 0.995, *p* = 0.338; **Fig 3c)**. Similar to our previous study (Margetts-Smith *et al*., 2021), EE reduced Fos (*t*_(16)_ = 3.496, *p* <0.01; **Fig 3d)**. Specifically, Fos expression in PL→PVT neurons was significantly lower in the EE compared to SH condition (*t*_(13)_ = 2.729, *p* < 0.05), whereas there were no significant differences in Fos in PL→NAc neurons between EE and SH conditions (*t*_(15)_ = 0.042, *p* = 0.967; **Fig 3e)**. Finally, the percentage of PL→PVT neurons expressing Fos was significantly lower in the EE compared to SH condition (*U =* 2.0, p < 0.01), but no differences in the percentage of PL→NAc neurons expressing Fos between EE and SH conditions were observed (*U =* 34.5, *p* = 0.981, **Fig 3f)**

**Figure 3.**
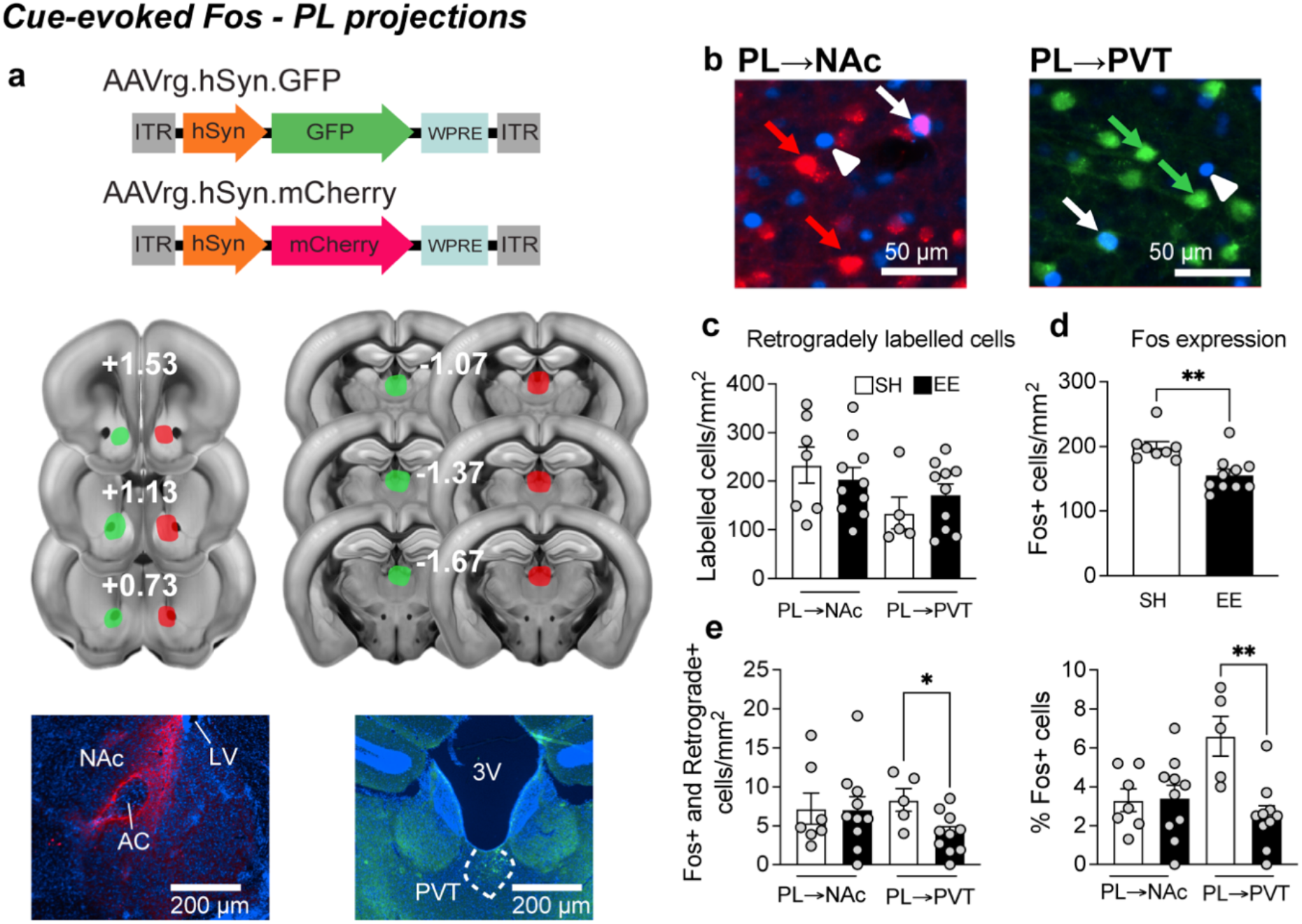
EE attenuated cue-evoked Fos in PL→PVT and interneurons, but not in PL→NAc neurons. **a** Overview of the retrograde AAVs used to express GFP and mCherry in the PL and their representative injection sites. **b** Representative images of Fos-expressing cells (white arrowhead), PL→NAc (red arrow) and PL→PVT (green arrow) neurons that co-express Fos (white arrow) **c** Similar levels of retrogradely labelled PL→NAc and PL→PVT neurons. **d** EE generally reduces Fos expression and **e** in PL→PVT, but not PL→NAc neurons. **f** Proportion of PL→NAc and PL→PVT neurons that express Fos. (c-f; PL→NAc (n=7-10/group) and PL→PVT (n=5-10/group). All data are expressed as mean±SEM.

## Discussion

Here we investigated how EE attenuated cue-evoked sucrose seeking via modulation of PL interneuron excitability and pyramidal cell recruitment. EE upregulated the baseline excitability within interneurons, but not pyramidal cells. Additionally, EE reduced recruitment of PL→PVT, but not PL→NAc, projection neurons. Our findings provide deeper insight into modification of PL circuits relevant for ‘anti-food seeking’, consisting of PL interneuron excitability overdrive and corticothalamic underdrive. We discuss below the EE-mediated neuronal adaptations and their implications on modulating cue-evoked sucrose seeking and neuronal recruitment and activity patterns.

### EE-induced enhancement of interneuron, but not pyramidal cell, excitability

EE increased interneuron excitability via enhancing its firing capacity; thereby increasing PL’s ability to exert more internal inhibitory control. Also, associated with this enhancement were decreased AP half-width and peak and increased mAHP and fAHP components of the action potential. Since these components are influenced by voltage-gated A-type K^+^ channels and Ca^2+^–dependent K^+^ channels (BK channels for fAHP and SK channels for mAHP), these alterations may reflect an alteration in function and/or number of these channels (Ishikawa *et al*., 2003; Villalobos *et al*., 2004; Kasten *et al*., 2007; Johnston *et al*., 2010; Wang *et al*., 2016). In general, K^+^ channels are associated with reducing excitability, but paradoxically, K^+^ channels can contribute to enhancing neuronal firing. For instance, the Kv3.3-3.4 subfamily of K^+^ channels that conduct A-type K^+^ currents enhance firing rates by rapidly de-inactivating voltage-gated Na+ channels, enabling faster recovery to resting conditions that allow subsequent neuronal firing (Kasten *et al*., 2007; Johnston *et al*., 2010; Jaffe & Brenner, 2018). BK channels have also been implicated in this increased firing effect through their interaction with e.g. L-type Ca^2+^ channels that enhance neuronal repolarization and decrease AP half-width (Wang *et al*., 2016). Further neurophysiological investigations are required to elucidate the precise intrinsic factors underlying these alterations in interneurons following EE.

One factor contributing to enhanced excitability could be the increased formation of perineuronal nets (PNNs), a network of extracellular matrix (ECM) molecules on interneurons. Indeed, PNNs enhance interneuron excitability (Balmer, 2016) and EE increases PNN intensity in mPFC interneurons (Slaker *et al*., 2016). PNNs typically surround parvalbumin-(PV) expressing interneurons, which inhibit the activity of nearby pyramidal cells and interneurons and facilitates the suppression of sucrose seeking (Sparta *et al*., 2014). Hence, it is possible that EE enhances PV+ interneuron excitability, which contributes to decreased Fos expression in PL→PVT neurons here. Thus, in future studies, it would be paramount to identify whether PV+ interneurons do indeed exhibit enhanced excitability, especially since we did not classify interneurons based on their phenotype. Indeed, several subtypes of interneurons exist, which are classified based on their neurochemical phenotypic markers such as parvalbumin (PV), somatostatin (SOM), and vasointestinal peptide (VIP) and exhibit differences in intrinsic physiology (Anastasiades & Carter, 2021).

Finally, we did not detect any changes in pyramidal cell excitability. One possibility for not detecting altered excitability is that we did not delineate between specific pyramidal cells, such as PL→PVT vs PL→NAc projection neurons that exhibited differential activity patterns following EE. Hence, subtle excitability changes may not have been detected. Hence, future investigations should separately examine these neuronal populations to provide clearer insights into EE-induced excitability adaptations.

EE attenuated recruitment of corticothalamic, but not corticoaccumbens, projections Related to the increased ability of PL interneurons to exert local inhibition, EE decreased recruitment of PL→PVT neurons. This decrease may reflect changes in encoding sucrose’s diminished value after EE, as evidenced by reduced sucrose consumption. Here, EE represents a much larger reward than sucrose and resembles aspects of a devaluation manipulation (Sieburg *et al*., 2019). Indeed, reward devaluation decreases the proportion of putative PL pyramidal cells excited by sucrose cues (Niedringhaus & West, 2022).

EE did not alter PL→NAc recruitment despite its role in reward seeking (Gourley & Taylor, 2016). However, this finding aligns with our previous study that EE did not decrease PL and NAc Fos expression (Margetts-Smith *et al*., 2021). Furthermore, PL ensembles tagged during sucrose operant conditioning send only minor projections to the NAc (Visser *et al*., 2020). Hence, EE may attenuate sucrose seeking via preferentially modulating PL→PVT recruitment and brain structures targeted by the PVT, such as the central amygdala that contains neuronal ensembles that suppress cue-evoked food seeking (Lay *et al*., 2023).

### Summary and future directions

Here we have extended our previous study (Margetts-Smith *et al*., 2021) by revealing how EE dynamically enhanced PL’s baseline interneuron excitability and attenuated recruitment of cue-evoked corticothalamic neurons. Future studies need to delineate the precise interneuron populations, and the downstream neuronal targets modulated due to this neuronal overdrive and underdrive. We highlight how experience-dependent plasticity within PL’s inhibitory circuits may contribute to reduced PL output and suppression of food seeking, thereby revealing PL interneurons and PVT-projecting neurons as a potentially critical component of the brain’s ‘anti-craving’ circuitry.

## Materials and methods

### Animals

C57BL/6J female mice (Charles River UK) aged 8-12 weeks at the beginning of experiments were group housed 2-4 per cage under a 12 h light/dark cycle (lights on at 7:00 A.M.) at a maintained temperature of 21 ± 1°C and 50 ± 5% relative humidity. One week before conditioning and until experiment completion, mice were food restricted to 90% of their *ad libitum* body weight. All experiments were conducted in accordance with the UK Animals (Scientific Procedures) Act 1986 (ASPA).

### Magazine Training and Pavlovian Conditioning

Similar apparatus and behavioral procedures were used as previously described (Ziminski *et al*., 2017; Brebner, Ziminski, Margetts-Smith, Sieburg, Reeve, *et al*., 2020). Briefly, behavioral training and testing were conducted in mouse conditioning chambers (15.9 × 14 × 12.7 cm; Med Associates, Vermont, USA). A syringe pump dispensed 10% sucrose solution (serving as the unconditioned stimulus (US) into a magazine receptacle. The conditioned stimulus (CS or ‘Cue’) was an auditory clicker. Med-PC IV (Med Associates) software controlled all experimental parameters and data collection.

Mice underwent a magazine training session followed by 10-12, twice daily conditioning sessions. Each session had 6 Cue presentations (120 second clicker) separated by RI-120s inter-trial intervals (No Cue period). During each Cue period, ∼15 µl deliveries of 10% sucrose solution were delivered on a RI-30 s schedule.

At 3-7 d following the last acquisition session, mice were exposed to 1d EE or left in standard housing (SH). In Experiments 1 and 3 mice then underwent a single test session under extinction conditions to assess cue-evoked food seeking. In Experiment 2, mice were immediately assessed for neuronal excitability following 1d EE.

### Environmental enrichment (EE)

Standard housing (SH) consisted of a small standard cage (15.9 × 14 × 12.7 cm) with basic nesting material and a wooden chew block. Environmental enrichment (EE) housing consisted of larger cages with 3 tiers (40 × 26 × 53 cm), with connecting tunnels, a separate sleeping pod, two exercise wheels, multiple nesting and bedding material, plastic houses, tunnels, wooden chew bars and Lego bricks **(Fig 1a**, (Margetts-Smith *et al*., 2021).

### Virus information

#### Adeno-associated viruses

**Experiment 2**– For expression of mRuby in inhibitory neurons we used AAVDJ/8/2-mDlx-mRuby (viral titer 6.9 × 10^12^ vg/ml; catalog # v242-DJ/8, Viral Vector Facility (VVF) of the Neuroscience Center Zurich (ZNZ), constructed using elements from mDlx-HBB-chI: Addgene #83900 and mRuby3: Addgene #85146) we diluted this virus 1/10 with sterile saline.

**Experiment 4** – For retrograde AAV tracing; AAVrg-hSyn-EGFP (viral titer 7.0 × 10^12^ vg/ml; catalog # 50465-AAVrg, Addgene, Deposited by Bryan Roth) and AAVrg-hSyn-mCherry (viral titer 7.0 × 10^12^ vg/ml; catalog # 114472-AAVrg, Addgene, Deposited by Karl Deisseroth) diluted 1/10 with sterile saline.

### Surgical procedures

Mice (between 8-11 weeks old) were anaesthetized with isoflurane, mounted onto a stereotaxic frame (RWD Life Science) and received pre-operative analgesia (s.c Meloxicam). An approximately 1cm incision was made with a scalpel and hole(s) was drilled above the area(s) of infusion. A sterile pulled glass pipette (Drummond PCR Micropipets 1-5 µl) was attached to a syringe pump (World Precision Instruments, UMP3 micropump) and then filled with virus. Viral infusion pipette was slowly lowered to the desired DV co-ordinate and 500 nl of virus was injected at 100 nl per minute (300 nl in the case of retrograde tracing experiments). For mRuby expression to label inhibitory cells, bilateral injections were made in the PL (AP +1.8 mm, ML ± 0.35 mm, DV: -2.4 mm relative to bregma). For retrograde tracing experiments using retrograde-GFP and retrograde-mCherry viruses these were injected unilaterally into the nucleus accumbens (NAc: AP +1.2 mm, ML ± 1.1 mm, DV: -3.9 mm from brain surface) and paraventricular nucleus of the thalamus (PVT: 20° angle, AP - 1.36 mm, ML ± 1.13 mm, DV: -3.3 mm relative to bregma). Mice were left 3-5 weeks for recovery and viral expression before experiments began.

### Immunohistochemistry and Fos quantification

Fos staining was performed as previously described (Ziminski *et al*., 2017; Brebner, Ziminski, Margetts-Smith, Sieburg, Reeve, *et al*., 2020). Briefly, ninety minutes following initiation of the final test session mice were anaesthetized and then transcardially perfused with 4% PFA, then cryoprotected with 30% sucrose in PBS 1X before being frozen on dry ice and stored at -80°C. Coronal sections (30 µm) were taken using a Leica CM1900 cryostat and contained the PL (corresponding to approximately AP 2.13-1.73; (Paxinos & Franklin, 2001). Whole brains were sectioned to include NAc and PVT injection sites.

Sections were incubated at RT overnight in 1/300 anti-Fos primary antibody (cat# 2250, RRID:AB_2247211; Cell Signaling Technology), and for tracing experiments 1:1000 anti-GFP (catalog #ab13970, RRID:AB_300798; Abcam). The following day, slices were washed in PBS then incubated in their corresponding secondary antibodies for 90 minutes. Secondaries used at 1/300 dilution were: anti-rabbit anti-rabbit Alexa 647 (for Fos detection, catalog #A-21245, Thermo Fisher Scientific, RRID:AB_2535813), anti-chicken 568 (for GFP detection, catalog #SAB4600039, RRID: AB_2631230; Sigma-Aldrich). The fluorescent dye DAPI (catalog #D-9542, Sigma-Aldrich) was used for nuclear staining at 5 µM concentration. Slices were mounted on microscope slides (catalog # 11562203; Fisher UK) air-dried, and coverslipped with Fluoromount G (catalog #00-4958, RRID:SCR_015961, Invitrogen).

Fluorescence images of Fos staining, GFP and mCherry (Exp 3) from left and right hemispheres of the PL from 1–2 coronal sections per animal, corresponding to approximately bregma 2.13 to 1.73 (Paxinos & Franklin, 2001), were captured using a 10x objective lens (N.A. 0.3; Olympus) with a QI click camera (Qimaging) attached to an Olympus Bx53 microscope. Fos+ nuclei were quantified using iVision software (version 4.0.15, RRID: SCR_014786; Biovision Technologies). Images of Fos immunoreactive (IR) nuclei in the PL cortex were digitally merged into a single image from 10 Z-stacks using iVision, at multiple focal planes (10 µm thickness). Images were normalized in iVision by adjusting the contrast for each image to match the contrast function to the integrated histogram to a set range of values. Cell counts were performed blind. Representative images of regions of interest (ROIs) for cell counts were analyzed using iVision software (version 4.0.15, RRID: SCR_014786; Biovision Technologies).

### *Ex vivo* electrophysiology

#### Brain slice preparation

Slice electrophysiology was performed as previously described (Ziminski *et al*., 2017; Brebner, Ziminski, Margetts-Smith, Sieburg, Reeve, *et al*., 2020). One to two weeks from the final acquisition session, mice were exposed to 1d EE (or no EE in control mice), and their brains were rapidly removed and immersed into near-freezing oxygenated high-sucrose artificial cerebrospinal fluid (S-aCSF) with the following composition (mM): 60 sucrose, 87 NaCl, 2.5 KCl, 3 MgCl_2_, 0.5 CaCl_2_, 26 NaHCO_3_, 1.25 NaH_2_PO_4_ H_2_0, and 10 D-glucose, saturated with 95% O_2_ /5% CO_2_. Coronal slices containing the prelimbic cortex (PL) of the mPFC. (250-300 μm thick sections; ∼bregma 1.5-2.4 mm) were sliced using a Leica VT1200S slicer, stored in a chamber containing carbogenated-aCSF at ∼34°C for 30 minutes, then cooled to room temperature for a further 10-30 minutes prior to recordings. For recordings, slices were transferred to a recording chamber and perfused at 2–3 ml/min with 30–32°C standard aCSF containing (mM): 125 NaCl, 2.5 KCl, 2 CaCl_2_, 1 MgCl_2_, 26 NaHCO_3_, 1 NaH_2_PO_4_ H_2_0, 10 D-glucose, and bubbled with 95% O_2_ /5% CO_2_.

#### Whole-cell recordings

Whole-cell recordings on PL layers V-VI pyramidal cells or interneurons in Experiment 2 **(Fig. 2)** were performed using glass-pipettes (1.5 mm outer diameter, 0.86 mm inner diameter) for intrinsic excitability recordings. Intracellular solutions consisted of (in mM): 125 K-gluconate, 10 KCl, 2 MgCl_2_, 0.1 CaCl_2_, 10 HEPES, 1 EGTA, 2 Mg-ATP and 0.2 Na-GTP (pH 7.2-7.4). Pyramidal cells were identified based on their morphology and/or characteristic firing properties similar to previous studies (Cao *et al*., 2009; Ziminski *et al*., 2017; Brebner, Ziminski, Margetts-Smith, Sieburg, Reeve, *et al*., 2020). Interneurons were identified based on their mRuby expression with a 561 nm excitation wavelength.

Data were collected with a Multiclamp 700B amplifier (Molecular Devices), A/D board (PCI 6024E; National Instruments) and WinWCP Software (courtesy of Dr. John Dempster, University of Strathclyde, Glasgow, UK. John Dempster (http://spider.science.strath.ac.uk/sipbs/software_ses.htm). Signals were amplified, filtered at 4 kHz and digitized at 10 kHz. The Hum Bug noise eliminator (Quest Scientific) was used to reduce noise. Neurons were visualized with differential interference contrast using an Olympus BX51WI microscope attached to a Revolution XD spinning disk confocal system (252, Andor Technology) for fluorescence microscopy.

#### Intrinsic excitability recordings

Pyramidal cells and interneurons were held at −65 mV and -80 mV, respectively, for the recording duration. The current clamp protocol consisted of 800 ms current injections from −60 pA to 730 pA incrementing in 10 pA steps for pyramidal cells and from -60 pA to 450 pA incrementing in 5 pA steps for interneurons. The liquid junction potential was −13.7 mV and was unaccounted for. Spike counts, spike kinetics and input resistance were analyzed with Easy Electrophysiology software (Easy Electrophysiology Ltd).

## Data Analysis

All statistical tests were performed using Prism software (RRID:SCR_002798; GraphPad 10 Software). For all experiments, data was analyzed using multifactorial ANOVAs, followed by post-hoc testing or pair-wise comparisons using a 2-tailed t-test. Group data are presented as mean ± SEM. Data points exceeding +/-2 SDs from the mean were excluded as outliers.

## Acknowledgments

We would like to thank Dr. Andre Chagas and Prof. Hans Crombag (University of Sussex), Dr. Alex Hoffman (NIDA/NIH), Dr. Gabriella Margetts-Smith (University of Bristol), Lauren French and Lucia Scasso for technical support and/or discussions. We also thank Dr. Joseph Ziminski (Sainsbury Wellcome Center) for technical support. We thank Drs Bryan Roth, Karl Deisseroth and James M Wilson for gifting their plasmids to Addgene and allowing us to purchase and use them via Addgene. We acknowledge the Viral Vector Facility (VVF) of the Neuroscience Centre Zurich for providing the plasmids used in this study.

